# Goal-Directed Learning in Cortical Organoids

**DOI:** 10.1101/2024.12.07.627350

**Authors:** Ash Robbins, Hunter E. Schweiger, Sebastian Hernandez, Alex Spaeth, Kateryna Voitiuk, David F. Parks, Tjitse van der Molen, Jinghui Geng, Tal Sharf, Mohammed A. Mostajo-Radji, David Haussler, Mircea Teodorescu

**Affiliations:** Department of Electrical and Computer Engineering, University of California Santa Cruz, Santa Cruz, CA 95064, USA; Genomics Institute, University of California Santa Cruz, Santa Cruz, CA 95064, USA; Department of Molecular, Cell, and Developmental Biology, University of California Santa Cruz, Santa Cruz, CA 95064, USA; Department of Biomolecular Engineering, University of California Santa Cruz, Santa Cruz, CA 95064, USA; Neuroscience Research Institute, University of California Santa Barbara, Santa Barbara, CA 93106, USA; Department of Molecular, Cellular and Developmental Biology, University of California Santa Barbara, Santa Barbara, CA 93106, USA

**Keywords:** biocomputing, organoid intelligence, artificial intelligence, reinforcement learning, electrophysiology, stimulation, electrical stimulation, in vitro, neurons

## Abstract

Experimental neuroscience techniques are advancing rapidly, with major recent developments in high-density electrophysiology and targeted electrical stimulation. In combination with these techniques, cortical organoids derived from pluripotent stem cells show great promise as *in vitro* models of brain development and function. Although sensory input is vital to neurodevelopment *in vivo*, few studies have explored the effect of meaningful input to *in vitro* neural cultures over time. In this work, we demonstrate the first example of goal-directed learning in brain organoids. We developed a closed-loop electrophysiology framework to embody mouse cortical organoids into a simulated dynamical task (the inverted pendulum problem known as ‘Cartpole’) and evaluate learning through high-frequency training signals. Longitudinal experiments enabled by this framework illuminate how different methods of selecting training signals enable improvement on the tasks. We found that for most organoids, training signals chosen by artificial reinforcement learning yield better performance on the task than randomly chosen training signals or the absence of a training signal. This systematic approach to studying learning mechanisms *in vitro* opens new possibilities for therapeutic interventions and biological computation.

## Introduction

Neurons have an unmatched capability for intelligent and dynamic information processing. Deep artificial neural networks require over five layers to achieve an adequate approximation of a single biological neuron [1]. Modern methods of interfacing with neural tissue involve any combination of encoding information, decoding information, or perturbing the underlying dynamics through various timescales of plasticity [2–6]. These biological networks operate at a fraction of the power consumption of artificial systems [7]. Understanding how they process and dynamically adapt to information has profound implications, from developing targeted stimulation protocols for neural rehabilitation to creating energy-efficient bio-electronic computation [8].

*In vivo* training methods commonly exploit principles of reinforcement learning [9–11] and Hebbian learning [2, 12, 13] to modify biological networks. However, *in vitro* training has not seen comparable success, and often cannot utilize the underlying, multi-regional circuits enabling dopaminergic learning [14]. Early work by Bakkum et al. [15]) demonstrated promising results, yet there have been few significant advances in *in vitro* learning since then. Our knowledge of biological learning rules has not yet translated to reliable methods for consistently training neural tissue in a goal-directed fashion. Successfully harnessing *in vitro* learning could reveal fundamental mesoscale learning principles. We developed a training paradigm that applies fundamental training patterns to achieve control in a dynamical task by embodying an organoid in a virtual environment.

Pluripotent stem cells-derived brain organoids have been shown to recapitulate key aspects of neuronal development, including diverse cell types and complex electrophysiological circuitry [16, 17]. Although these organoids can model many aspects of development, they are typically grown in mono-directed differentiation, lacking key inputs from other regions [18, 19]. The combination of organoid complexity with high-density microelectrode arrays (HD-MEAs) enables precision in both recording and stimulating at single-cell resolution [20]. This fine-grained control allows us to systematically perturb these networks and longitudinally observe their responses, providing a platform for iterative investigation of learning at the circuit-level.

Within the last three decades, labs have used varying methodologies of stimulation in efforts to harness biological computation through closed-loop interaction[15, 21, 22]. Pioneering work demonstrated this by repurposing vestibular circuits of a lamprey to control a robot [23], leading to numerous studies embodying cultures into closed-loop tasks [24–31]. Different frameworks for *in vitro* learning have emerged, influenced by factors like species, brain regions, neuron distributions, and network architectures. Early work accomplished supervised learning through slow frequency stimulation which evoked network bursts, ceasing stimulations when the desired effect was achieved [32, 33], formalized as learning by stimulation avoidance [34]. Many approaches use tetanic (high-frequency) stimulation patterns as training signals [5, 15]. These tetanic electrical stimulations (*>* 5 Hz) have been shown to induce functional and stimulus-evoked changes for pattern recognition [35], burst information [5], exploring LTP/LTD [36–39], and embodied tasks [25, 26, 28]. Tetanic stimulation is thought to induce changes through associative mechanisms due to increased neural activity. Reservoir computing, which leverages machine learning readout layers, has demonstrated success in unsupervised tasks like predicting dynamical equations and classifying speech snippets [22], as well as a control task [30]. The free energy principle has been proposed as a theoretical framework to explain blind source separation tasks [40, 41] and control in a simplified “Pong” environment [21], drawing parallels to artificial reinforcement learning approaches to curiosity [42]. Recent work has shown that even inanimate hydrogels, when electrically embodied in a “Pong” task space, can achieve learning [43], highlighting the importance of appropriate controls and careful interpretation and investigation of apparent learning in these systems. While previous work has demonstrated that stimulation can modify synaptic plasticity, key questions remain about the optimal selection of target neurons, stimulation frequencies, and timing.

In this paper, we propose a systematic approach to characterizing and training biological controllers through high-frequency training signals within a task-based framework. First, we employ automated analysis to yield putative neurons, units which we then characterize through repeated electrical stimulations. Next, we derive causal connectivity metrics from the stimulation responses, which are used to determine a neural configuration. Finally, we embody the cortical organoids in the classical inverted pendulum control stabilization problem [44, 45] (similar to balancing a ruler vertically on your hand). We used a two dimensional “virtual” cart that can move on the horizontal axis to balance a vertical pole (Fig. 1). This is commonly known as “cartpole” [46, 47]. Unlike pattern recognition tasks, this problem requires continuous, active stabilization of an inherently unstable system where small lapses in control lead to failure - making it particularly suitable for assessing both real-time processing and adaptation. We use short, high-frequency training signals, and investigate three distinct methods of delivering training patterns: “*Null*”, “*Random*”, and “*Adaptive*”, demonstrating that adaptive training significantly outperforms the alternatives. Through extended experiments with continuous adaptive training, we further validate this approach while showing how our proposed causal connectivity metrics correlate strongly with performance outcomes.

**Fig. 1.**
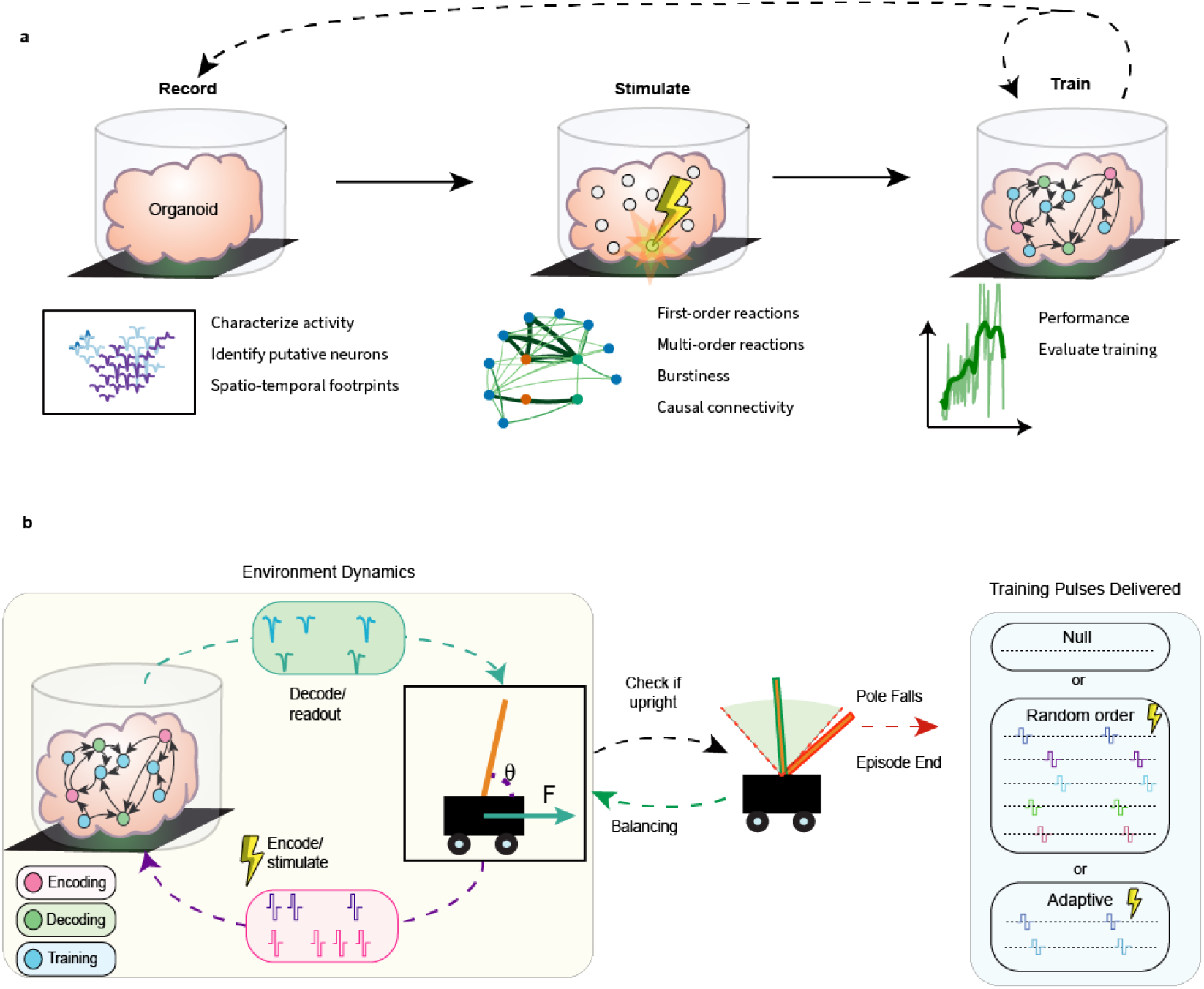
Experimental overview for *in vitro* learning. **a** Schematic of multiphase experimental design. The *Record* phase uses a spontaneous recording to locate and characterize putative neural units. The *Stimulate* phase uses electrical stimulation on each of these units to measure stimulus-evoked activity through different temporal ranges. Human experimenters then select putative neuron roles from the causal connectivity analysis. The *Train* phase consists of repeated interactions with the simulated dynamical environment, organized into episodes. **b** Flowchart of each episode of the training loop. The virtual environment is simulated at a fixed frame rate while interacting with the organoid in a closed loop (yellow box). The episode is terminated when the pole falls into an unrecoverable position. Finally, depending on the training paradigm, a pattern of pulses may be delivered.

## Results

### Multi-Phase Experimentation

We developed a framework that consists of three key experimental phases to achieve goal-directed learning in cortical organoids: 1) network characterization through spontaneous recording, 2) stimulus-response mapping through targeted electrical stimulation, and 3) closed-loop training in the dynamical task (Fig. 1a). Each phase builds upon automated analysis from the previous phase to systematically identify and interface with relevant neural circuits. This approach enables us to first characterize the network’s causal connectivity before attempting to dynamically modify the stimulus-evoked responses through training. The framework provides millisecond-precision control to minimize latency between the neural culture and virtual environment, and it also supports reproducibility through an automated analysis pipeline.

To implement this framework, we generated mouse cortical organoids to serve as a biological substrate for learning. Functional neural networks were developed through directed patterning and self-organization (Fig. 2a) of three-dimensional aggregates from embryonic stem cells into structured neural tissue (Fig. 2b) that recapitulates key features of cortical architecture [48]. By day 10, the networks show forebrain specified radial glia (Pax6) and medial ganglionic eminence progenitors (Nkx2.1), maturing by day 30 to express subtype-specific markers including upper (Satb2) and deep (Tbr1) layer neurons (Fig. 2c) along with inhibitory neurons (Sst) and astrocytes (Gfap) Supplementary Fig. S2. We specifically chose cortical patterning due to the cortex’s well-established role in adaptive information processing and its capability to encode, decode, and modify responses to novel inputs [49]. The organoids were interfaced with high-density microelectrode arrays (HD-MEAs) [50–52] (Fig. 2d-f), providing precise spatio-temporal control over the culture with a high number of putative neuronal units for potential computation.

**Fig. 2.**
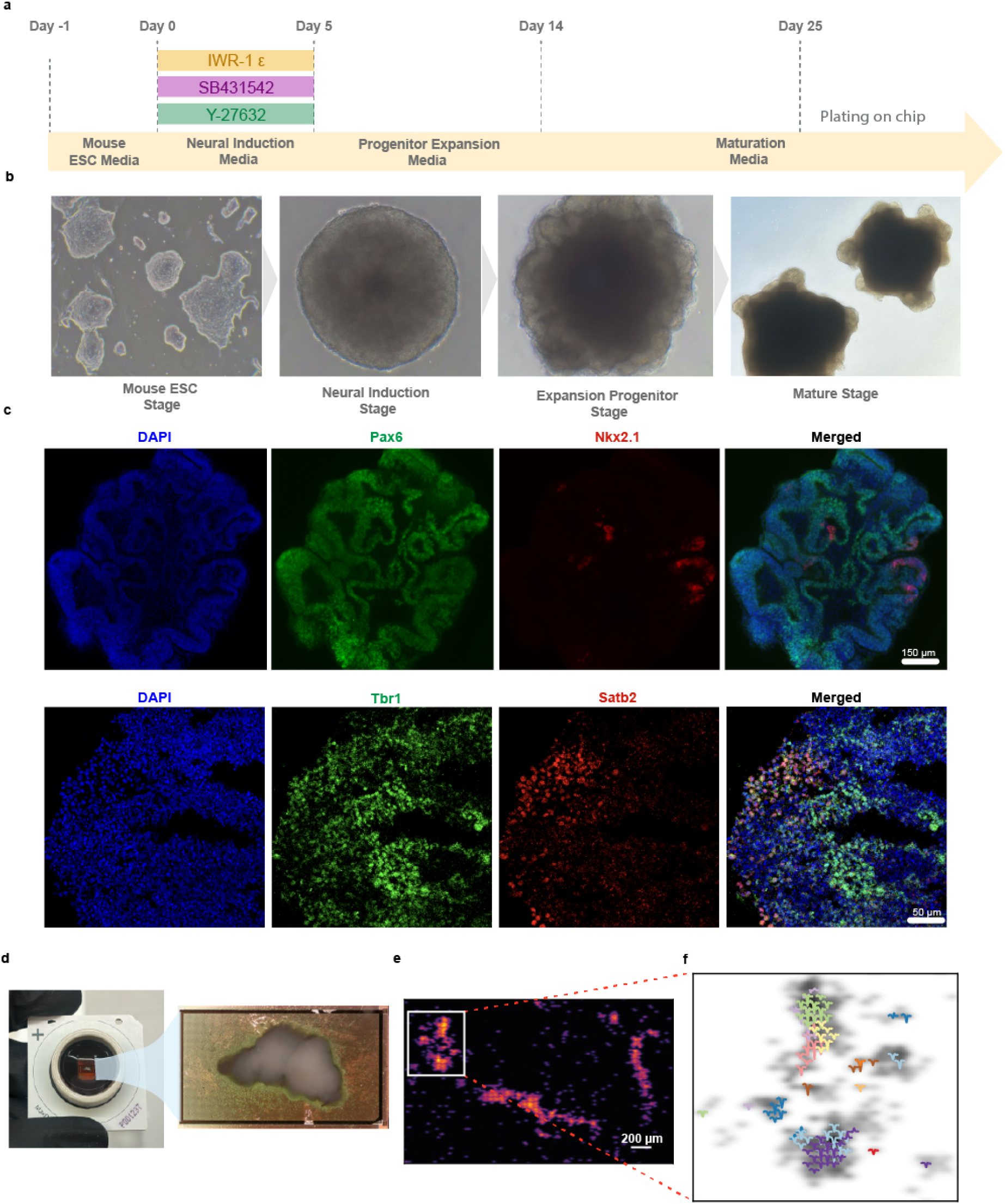
Organoid generation and interaction with electrical interface. **a** Schematic of the protocol for generating mouse embryonic stem cell (ESC) derived cortical organoids. **b** Brightfield images of organoids over development. **c** Immunohistochemistry of day 10 cortical organoids show the presence of Pax6 (radial glia), Nkx2.1 (medial ganglionic eminence) and Day 30 organoids express post-mitotic excitatory neuronal subtypes Tbr1 (deep layer) and Satb2 (upper layer). **d** Illustration of organoid plated on HD-MEA chip. **e** Heatmap of activity from electrode configuration overlayed on the HD-MEA and a representative plot of the electrode configuration. **f** Representative plot of the waveforms from individual neuronal units overlayed on electrode configuration

### Neural Configuration

Our targeted approach to cortical organoid computation focuses on characterizing capabilities within small sub-circuits. To do this, we require a few separate components. First, we begin by capturing spontaneous neural activity in the *Record* phase, using it to identify locations of putative neurons and the best electrodes to stimulate them. As shown in a previous study, electrically stimulating the axon initial segment yields the best chance at triggering action potentials [53]. The spontaneous activity recording is used to generate a spatial map of putative neural unit locations through a metric (Methods: Record Phase) incorporating firing rate and action potential amplitude (Fig. 2e,f). We then perform signal-averaging triggered by the local maxima at these locations to extract spatiotemporal footprints and remove duplicates for each unit. Larger amplitudes yield neurons which are easier to identify in real-time experiments. This analysis enables us to reliably detect neural activity on distinct electrodes throughout the experiment.

To map causal information flow through these neural circuits, we characterized stimulus-response relationships during the *Stimulate* phase by delivering bi-phasic pulses to each identified neural unit (50 pulses at 2Hz) as seen in Fig. 3a and c. Automated analysis quantified three major temporal response signatures: first-order causal connectivity representing direct neural pathways (Fig. 3b), multi-order causal connectivity showing network-mediated responses (Fig. 3d), and probability of evoking network-wide bursts (Supplementary Fig. S3). Based on these metrics, we created a neural configuration defining units for encoding, decoding, and training. We select two encoding and two decoding units, prioritizing pairs with strong first-order causal connectivity to maximize information transmission potential, and used multi-order connectivity as a secondary preference (Fig. 3e and f). Units exhibiting frequent network bursting responses were deemed less suitable as encoding units since widespread activation could interfere with more fine-grained control. After encoding and decoding units were selected, between 5–12 training units were selected independent of connectivity patterns to explore optimal training stimulations.

**Fig. 3.**
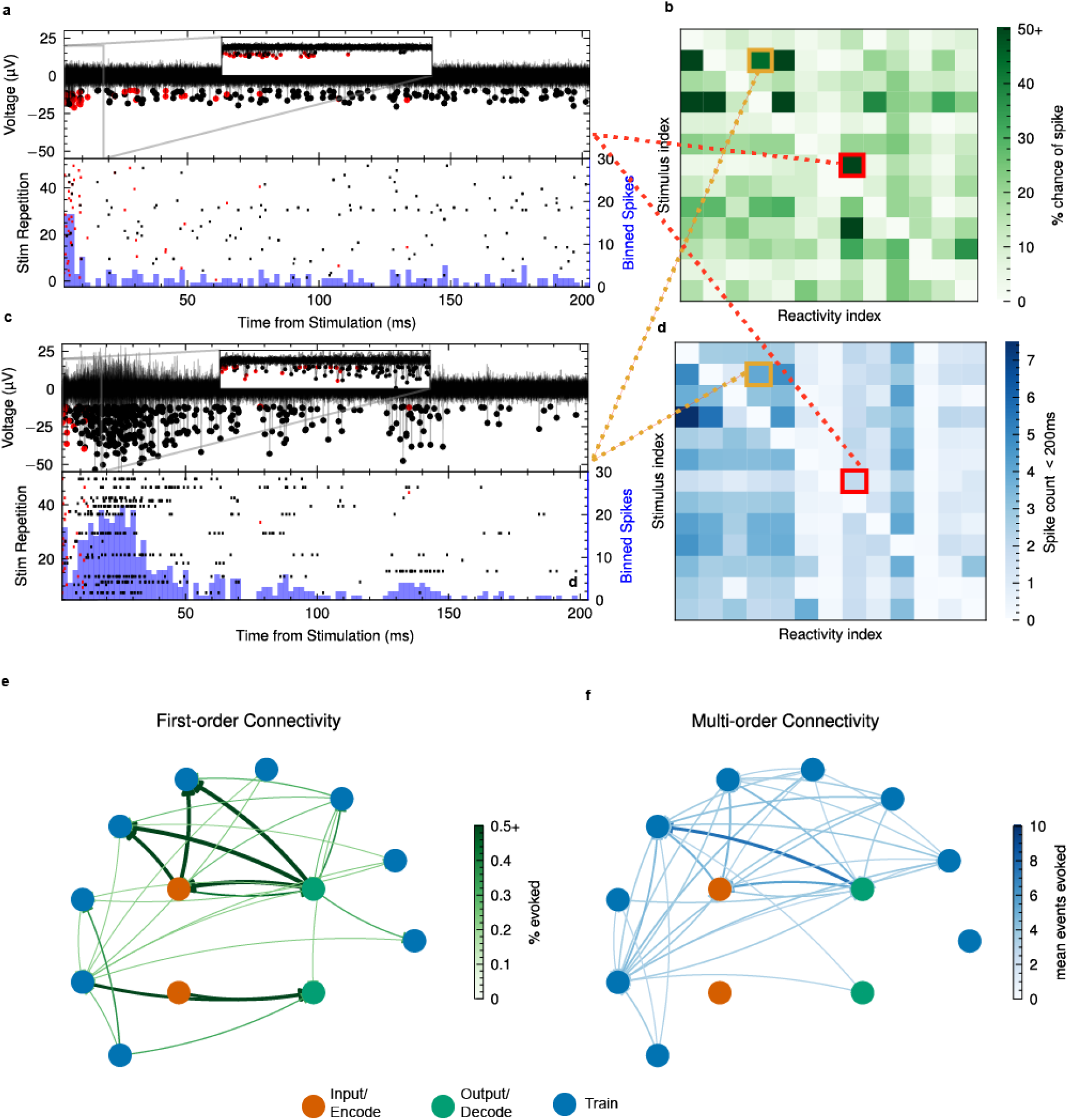
Electrical stimulation characterization and causal connectivity. **a** Stimulus-evoked responses showing (top) overlapping voltage responses from multiple stimulation repetitions and (bottom) latency from stimulation raster combined with peri-stimulus time histograms. This example highlights short-term responses. Inset shows the first 20ms. **b** First-order causal connectivity heatmap displaying the probability that a stimulus input evokes a reaction event within 18ms for the corresponding electrodes of interest. **c** Similar to a, highlighting a multi-order bursting response. **d** Similar to b, showing mean response count within 200ms. **e** First-order causal connectivity with chosen roles for an experiment. **f** Similar to e however with multi-order causal connectivity.

### Training Paradigm

Using our characterized neural configuration, in the *Train* phase we trained organoids to balance a pole on the cartpole task, where an inverted pendulum must be balanced by applying horizontal forces (between −10 N and 10 N) to the cart (Fig. 1). We implemented a rate-coding scheme [11, 54, 55] for both encoding pole angle and decoding motor commands as detailed in Methods: Encoding and Decoding Information. Each learning episode ran from initialization until the pole exceeded a terminal angle of ±16^*°*,^ which generally represents an unrecoverable state. A series of episodes is called a training cycle. To allow for both learning and recovery, experiments were organized into typically 15-minute training cycles with 45-minute rest periods. Performance-based feedback was delivered to the training units only at the ends of episodes when the organoid failed to balance the pole; no training feedback was given during the time it was balancing. Specifically, training signals were administered only when the 5-episode mean performance fell below the 20-episode moving average, allowing the system to focus on larger-scale adaptations while filtering out the impact of individual episodes of poor performance. If the time balanced never surpassed 10 s within the first cycle, different encoding/decoding units were selected. See Methods: Cartpole Environment for further details on training.

### Superior Performance of Adaptive Training Signals over Random and Null Conditions

To investigate whether training signals sent to the training neurons could modify information processing between input and output neurons, we compared three distinct experimental conditions: 1) “*Null*” without stimulation, 2) “*Random*” with stimulation of 5 randomly chosen training units (neurons), and 3) “*Adaptive*” with stimulation of a pair of training units (neurons) selected through reinforcement learning with eligibility traces - a technique that weights unit pairs based on their historical contribution to performance improvements (Fig. 4a,b). We refer to a 5-tuple or a pair of units selected for training stimulation feedback as a *training pattern*. Random-ordered 5-tuples were chosen to replicate previous studies [15, 21] and pairs were used adaptively as a more fundamental training signal. Refer to Methods Section: Training Signal Implementation for a detailed explanation of the training options tested. In each experiment, we tested two or more of these conditions in sequential 15-minute cycles (e.g., Null → Random → Adaptive) with one condition per cycle. Fig. 4c showcases a representative experiment where the adaptive condition (green) repeatedly achieved superior performance, improving from a baseline of 10 seconds to over 60 seconds of balanced control across multiple cycles. This improvement is quantified in the cycle-averaged performance metrics (Fig. 4d). The emergence of effective control behavior becomes evident when examining the pole angle trajectories over time (Fig. 4e). The effects of training stimulation delivered when the 5-episode mean performance dropped below the 20-episode mean is illustrated for three example cycles (Fig. 4f).

**Fig. 4.**
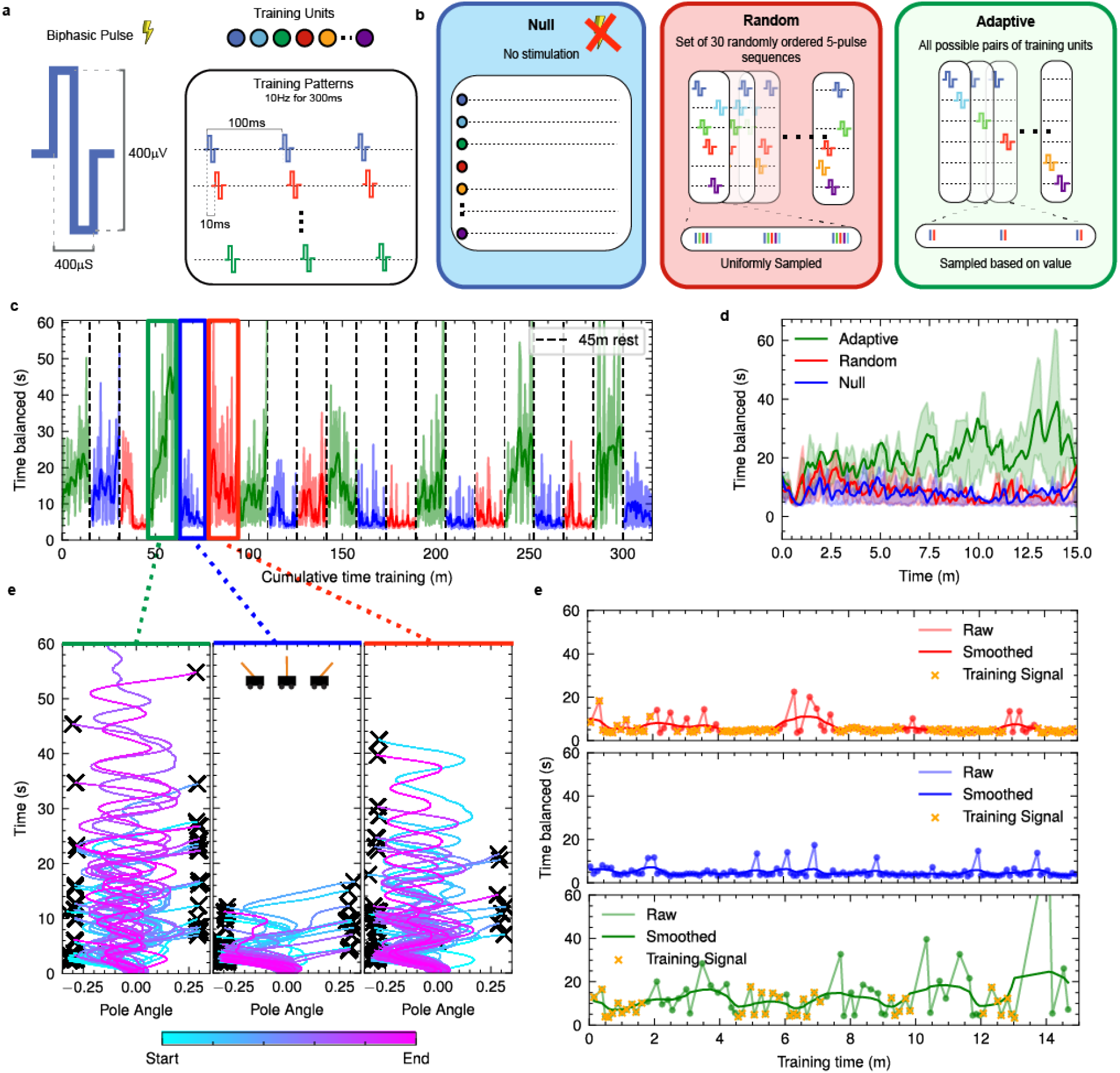
Training paradigms enable task-specific learning and performance evaluation. **a** Square wave biphasic pulse shape, 400µV peak to peak, 400µs period positive first. Training patterns consist of multiple pulses on separate channels spaced by 10ms within each pulse pattern, repeating the pattern at 100ms or 10Hz. **b** Description of the three separate training paradigms. Null: no stimulation, Random: 5-pulse patterns with the order randomly sampled from all possible training units, Adaptive: 2-pulse patterns chosen based on performance improvements attributed to specific stimulations. **c** A alternating experiment showing performance (time balanced in seconds) where the training paradigm is alternated, indicated by color (null-blue, red-random, green-adaptive). Each cycle lasts 15 minutes, with a 45 minute rest between cycles for 21 cumulative hours. **d** Mean and inter-quartile range of performance per training paradigm for the experiment in c. **e** Overlaid trajectories of pole angle throughout time within each training cycle for chosen cycles. **f** Plots of individual cycles, with each episode shown as scatter points, and training delivery times shown for the relevant episodes.

We evaluated 16 cortical organoids across 38 experiments, totaling over 125 hours of recorded activity. Organoids underwent 1–6 experiments each (median: 2) spanning 3–75 cycles (median: 27). The most extensive experiment ran continuously over 48 hours. To quantify consistent performance across experiments, we used the 90th percentile of episode duration (time balanced) within each cycle as our primary metric. We then established a rigorous benchmark for successful control based on the top 1% of performance achieved by the best-performing *in silico* random control algorithm (Supplementary Fig. S4). This threshold separates experiments where organoids reliably achieved proficient control from those showing only transient or inconsistent performance.

From these training paradigms (Null: *n* = 131, Random: *n* = 68, Adaptive: *n* = 92), adaptive stimulation significantly outperformed both random and null cases (*p* = 1.06e-3 and *p* = 2.45e-8 respectively, Mann-Whitney *U* test). Random stimulation outperformed the null case (*p* = 0.031), but to a lesser degree. Still, this shows that organoids can still learn something from a completely generic and randomly varying feedback signal when it is administered specifically when performance drops. While 22.8% of cycles reached proficiency under adaptive training, only 4.4% did so with random stimulation, and 2.3% with no stimulation.

Given the superior performance of adaptive training, we conducted extended experiments using only the adaptive paradigm without cycling between conditions. We will refer to this training strategy as “*Continuous Adaptive*” training. Fig. 5a demonstrates sustained learning across multiple hours, with performance consistently exceeding the proficiency threshold. The temporal pattern of performance peaks shows clear auto-correlation (Supplementary Fig. S1.2), suggesting underlying state-dependent changes occurring over multi-hour periods. Post-hoc analysis revealed certain pulse combinations yielded consistently higher improvement metrics (Fig. 5b), measured as the cumulative change in time balanced following each training signal delivery. These highly effective patterns—identified by improvement exceeding random walk bounds—often shared common training units. To elucidate how the eligibility trace-based value estimation method optimizes training signals, Fig. 5c visualizes the dynamic tracking of 2-unit training pattern value during an experiment with multiple cycles. The inset highlights how individual pulse patterns could drive either improvement or deterioration depending on the network’s state, emphasizing the importance of adaptive training signal selection.

**Fig. 5.**
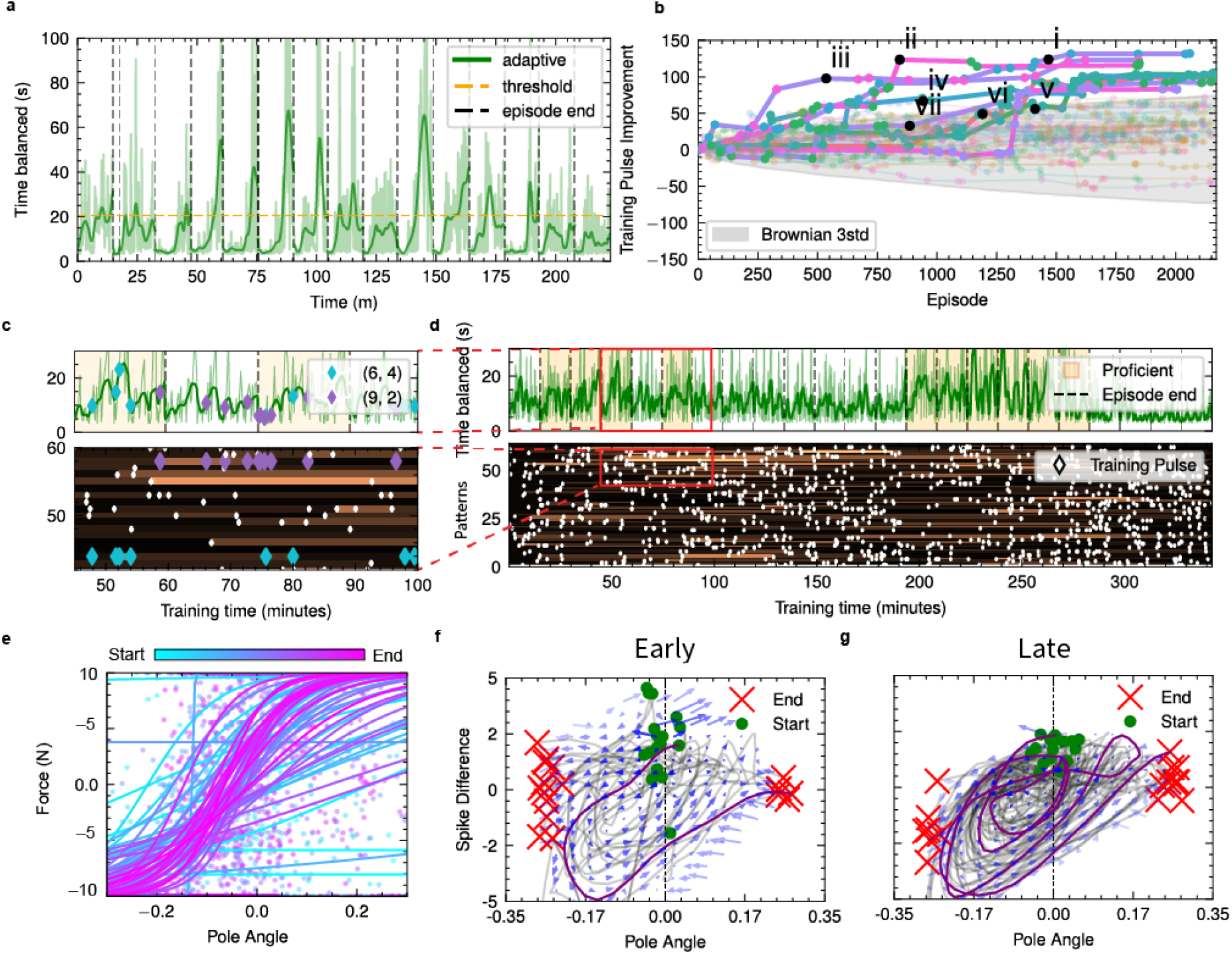
Continuous adaptive training reveals learning dynamics and control policy development. **a** Performance plot of the adaptive training paradigm running continuously for all cycles. Threshold indicates 20.5s which was designated as *proficient* Supplementary Fig. S4. **b** Improvement metric for various training patterns. Line color denotes first neuron in training pattern, scatter color denotes second neuron in training pattern. Grey shaded envelope designates brownian motion as explained in Supplementary Fig. S5. Roman numerals indicate highest maximum improvements. **c** Inset of performance through time with corresponding value estimation of training signals. The inset highlights two specific training patterns in blue and purple, showing their value changing through performance gain after the feedback stimulation pulses. With blue, a later pulse results in decreasing performance, thus the value of the blue pattern is decreased correspondingly. **d** Zoomed-out graph of previous inset, indicating longer-term value changes of performance through an entire experiment. **e** Sigmoid estimations of the organoids control policy through one training cycle. Early episodes show no cohesive structure, late episodes approach a sigmoid centered around 0°. **f** Early episodes (first third) dynamics of how the spike count difference between the output units respond to input frequencies dependent on the cartpoles angle. These responses are short and show less coherent flow patterns. **g** Similar to f, but late episodes (last third) show multiple round trips on the attractor and much higher density around a preferred state state near 0°. Late episodes also have less variability in initial responses.

We visualize a representative proficient organoid’s control strategy in two complementary ways. A simplified sigmoid policy estimation (Fig. 5d) shows the emergence of structured control centered around the vertical position, while the complete input-output flow fields in the phase space of neural activation patterns (Fig. 5f,g) reveal richer dynamics. These flow fields—which capture how neural responses relate to the pole’s state through time—show adaptation towards an off-center balancing point that accounts for both angle and angular velocity (see Supplementary Fig. S6). Early episodes show scattered, inconsistent responses, but late episodes demonstrate coherent control strategies with multiple stable oscillation patterns and increased activity density near this preferred state. In late episodes, the phase space of the dynamical system of pooled neural activations is also better at entraining the phase space trajectory from an initial preferred state into a 2-dimensional attractor manifold reflecting the problem’s control dynamics. A well-trained organoid circumnavigates this attractor many times.

These improvements in control strategy were reflected in overall performance metrics - continuous adaptive experiments achieved proficiency in 45.4% of cycles, substantially higher than the 22.8% rate observed in adaptive alternating experiments(Fig. 6a,b). We analyzed correlations between the best 90th percentile performance for each experiment and a range of connectivity metrics derived from both spontaneous and stimulus-evoked activity. Our first-order causal connectivity metric proved especially predictive of performance outcomes (*R*^2^ = 0.446, *p <* 0.001, *n* = 30), substantially outperforming traditional functional connectivity as calculated by spike-time tiling coefficient within 20 ms [56] (*R*^2^ = 0.288, *p <* 0.01, *n* = 32) (Fig. 6c,d). This advantage was most pronounced in proficient experiments, where causal connectivity showed remarkably strong correlation with learning performance (*R*^2^ = 0.58, *p <* 0.01, *n* = 12), whereas functional connectivity showed a weaker correlation (*R*^2^ = 0.200, *p* = 0.140, *n* = 12). The strength of first-order causal connections, which guided our neural configuration selection, emerged as a key predictor of learning capability (Fig. 6e). Output units’ ability to evoke multi-order responses (*p <* 0.01) and network-wide bursts (*p <* 0.01) correlated with performance, suggesting that output units’ capacity to recruit broader network activity may facilitate adaptive control. Other connectivity metrics, including burst probability and functional coupling between non-input/output units, showed weaker or non-significant correlations, further supporting the importance of first-order causal pathways in enabling successful learning.

**Fig. 6.**
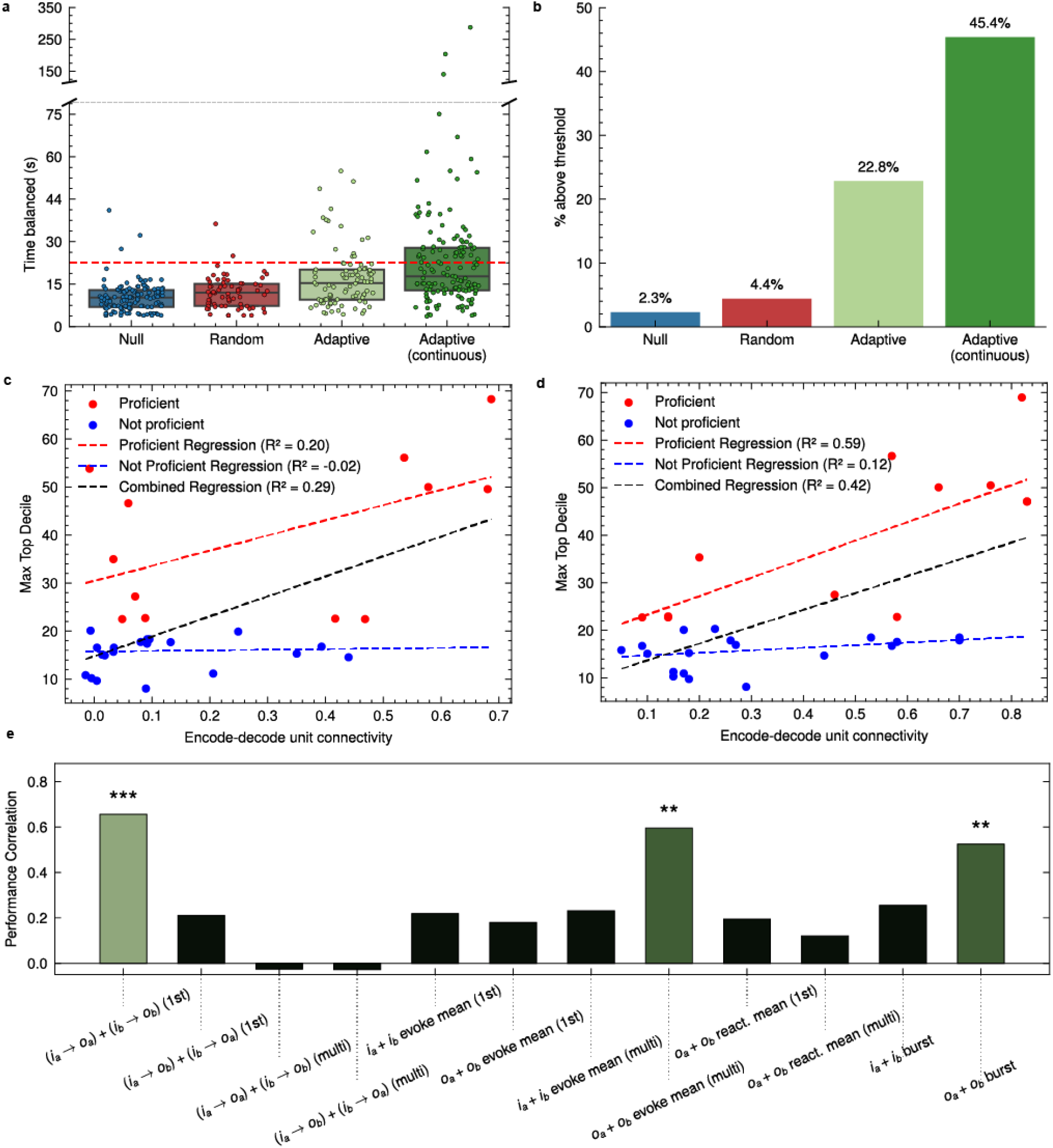
Statistical validation and neural correlates of learning performance. **a** Box plots of each condition with inter-quartile range where each datapoint represents the 90th percentile performance within a cycle (Null: *n* = 131, Random: *n* = 68, Adaptive: *n* = 92, Continuous Adaptive: *n* = 141 cycles), showing significant improvements (Mann-Whitney U test, *P* = 2.13 *×*10^−18^ vs null, *P* = 5.12 *×*10^−9^ vs random, *P* = 2.78 *×*10^−3^ vs alternating adaptive). We separate when adaptive was used for each cycle within an experiment versus when it was alternated. Threshold of “proficient” is shown as a red line. **b** Percentage of proficient cycles above the threshold. **c** Each experiments best 90th percentile performance predicted by functional connectivity calculated in the baseline recording (*R*^2^ = 0.288, *P* = 0.004, *n* = 32), along with regression lines fit to both proficient and not proficient cycles. **d** Similar to c, but with the first-order causal connectivity metric showing stronger predictive power (*R*^2^ = 0.446, *P* = 6.0 *×*10^−4^, *n* = 30). **e** Correlation of features with 90th percentile performance (*n* = 32). *i* and *o* represent the input and output units respectively.

These results demonstrate both the effectiveness of adaptive training and the importance of proper neural configuration selection. While high-frequency stimulation generally enhanced network performance over baseline, adaptive selection of training signals led to significantly better outcomes than random selection, with proficiency rates doubling in continuous experiments. The state-dependent nature of each cycle’s 90th percentile performance suggests persistent network states conducive to adaptation, rather than independent and isolated initializations. Our experimental design—delivering training pattern only at episode completion and selecting training units distinct from input/output units—isolates the learning effects from potential artifacts. The strong predictive power of first-order causal connectivity guided successful neural configuration selection, though learning remained configuration-dependent even within individual organoids.

## Discussion

Our results demonstrate the first instance of goal-directed learning in brain organoids using intentionally selected circuits, establishing a new paradigm for biological computation beyond the successful unsupervised reservoir learning approach demonstrated in [22]. Organoids naturally recapitulate neural development and, to some extent, circuit architecture in three-dimensional space, providing a highly parallel substrate for spatial computation. HD-MEAs enable millisecond-precision closed-loop control, allowing us to map specific computational roles to individual neurons despite capturing only a subset of the total network activity. Using a novel multi-phase experimental approach, we designed a rapid analysis pipeline that enables iterative experimentation, characterizing neural responses and adjusting parameters within minutes between experimental phases. This capability for systematic investigation mirrors the rapid prototyping that accelerated artificial neural network development. Mouse cortical organoids, with their quick development cycle[57, 58], provide an ideal testbed for this engineering approach to biological computation. This framework opens new possibilities for both fundamental neuroscience and hybrid bio-electronic systems.

Unlike previous *in vitro* learning studies focused on pattern recognition or fixed behaviors [15, 22], our framework tackles a continuously unstable control problem requiring active state maintenance, demonstrating that biological circuits can optimize complex dynamical systems even without canonical reward circuitry. Notably, even random stimulation improved performance over baseline, suggesting that high-frequency stimulation alone can modify network dynamics. However, our adaptive training paradigm, which optimized training signal selection based on performance changes, significantly outperformed both random and null conditions achieving proficiency in 45.1% of continuous cycles, demonstrating that biological neural networks can be systematically modified through precise electronic control. Our results show that task-specific adaptation can be achieved instead just through targeted 300 ms electrical stimulation patterns.

The effectiveness of training signals showed clear state-dependence, building on observations from [15] that different stimulation patterns succeed at different times. This suggests an interaction between training signals and recent activity history, providing an opportunity to explore fundamental stimulus-based learning rules at the circuit level. While mechanisms like spike-timing-dependent plasticity or short-term synaptic facilitation may underlie individual neuron-to-neuron modifications, how these local changes contribute to larger networks remains a key question for learning *in vitro*. We observed rapid modulation of control policies through these brief training cycles while also discovering temporal correlation in the network’s ability to achieve high performance over multi-hour periods, revealing distinct timescales of network modulation. Although our focus was on short-term modulation of network control policies, the temporal structure of network behavior evolved continuously over hours without clear directional trends. This suggests that future work should explore how to intentionally shape these longer-term dynamics to enhance both initial performance and learning capacity. Such adaptation across multiple timescales demonstrates a key advantage of biological systems for hybrid bio-electronic applications: while our training signals enabled rapid tuning of task-specific responses, the observed long-term state dependencies suggest potential for shaping underlying network dynamics to support and enhance these rapid adaptations. Training adaptations in different organoids lead to similar inverted pendumlum-dynamics-modeling low-dimensional attractors in the phase space of the organoid neurodynamics, even though the details of the neural connectivity in each organoid differ greatly. This reflects the flexibility of task-switching of mammalian cortex in vivo.

Causal connectivity analysis reveals how interfacing with the network influences learning capability, providing a framework for predicting and optimizing performance. The strong predictive power of first-order causal connectivity (*R*^2^ = 0.59 in proficient cycles) compared to traditional functional connectivity metrics demonstrates the importance of direct, stimulus-evoked pathways in architecting successful learning. The correlation between output units’ ability to evoke network-wide responses and performance suggests that effective neural interfaces benefit from neurons capable of recruiting broader network activity. This methodology of causal characterization could inform both therapeutic interventions and the design of future biological computing systems. By isolating fundamental stimulus-based functionality in simplified circuits, we can systematically identify and explore learning rules that may scale to complex bio-electronic systems.

Our open-source Python-based platform `BrainDance` (https://braingeneers.github.io/braindance) enables exploration of biological learning mechanisms through flexible and accessible experimental design, following the rapid iteration paradigm that accelerated machine learning development. We demonstrated success with rate coding and brief training patterns, however, the current technical limitations include planar HD-MEA recordings capturing only activity on the organoid side in contact with the HD-MEA, potential multi-unit activity due to single electrode thresholding, and manual expert-selection of neural configurations. More comprehensive investigations should examine performance across organoid development stages, regional specifications, and systematic methods for neural role assignment and real-time detection [59]. The current approach using predefined input/output units could be enhanced through latent space representations, and the integration of local field potentials may provide additional insight into network dynamics. Our simple eligibility trace-based value estimation method proved effective, yet more sophisticated reinforcement learning approaches could further optimize training signal selection based on information from the task and current estimated control policy. Future work should also investigate how different stimulation patterns, frequencies, and encoding schemes might improve learning outcomes. By systematically addressing these limitations while expanding our experimental toolkit, we can bring biological neural circuit investigation into an era of rapid and reproducible improvement, enabling deeper understanding and more effective utilization of their computational capabilities.

## Materials and Methods

### Electrophysiology experiments

Mouse cortical organoids were plated, as previously described by our group [57]. We plated mouse cortical organoids at day 25 on MaxOne high-density multielectrode arrays (Maxwell Biosystems). Prior to organoid plating, the multielectrode arrays were coated in 2 steps: First, we performed an overnight coating with 0.01% Poly-L-ornithine (Millipore Sigma # P4957) at 37°C overnight. Then washed the plates 3 times with PBS. We then performed an overnight coating with 5 µg*/*mL mouse Laminin (Fisher Scientific # CB40232) and 5 µg*/*mL human Fibronectin (Fisher Scientific # CB40008) at 37°C.

After coating, we placed the organoids on the chip and removed excess media. The organoids were then incubated at 37°C for 10 minutes to promote attachment. We then added pre-warmed neuronal maturation medi and changed the media every 2-3 days.

### High-Density MEA Recording

Extracellular signals were obtained through the MaxWell MaxOne system (MaxOne, Maxwell Biosystems) [60], using custom experimental software for the precise stimulation setup and timing [51]. Signals were recorded at a sampling rate of 20 kHz/channel with a 1 Hz hardware filter, for up to 1024 channels. At most 32 stimulation electrodes could be selected at a time. Cultures were maintained within incubators fulltime during recordings. Experiments were ran with the `BrainDance` (https://braingeneers.github.io/braindance) python library and longitudinally scheduled using an Internet of Things framework [61]

### Embryonic stem cell maintenance

All experiments were performed in the adapted BRUCE-4 mouse embryonic stem cell (ESC) line (Millipore Sigma #SF-CMTI-2). This line is derived from a male of the C57/BL6J mouse strain. Mycoplasma testing confirmed lack of contamination.

ESCs were maintained on Recombinant Human Protein Vitronectin (Thermo Fisher Scientific # A14700) coated plates using mESC maintenance media containing Glasgow Minimum Essential Medium (Thermo Fisher Scientific # 11710035), Embryonic Stem Cell-Qualified Fetal Bovine Serum (Thermo Fisher Scientific # 10439001), 0.1 mM MEM Non-Essential Amino Acids (Thermo Fisher Scientific # 11140050), 1 mM Sodium Pyruvate (Millipore Sigma # S8636), 2 mM Glutamax supplement (Thermo Fisher Scientific # 35050061), 0.1 mM 2-Mercaptoethanol (Millipore Sigma # M3148), and 0.05 mg*/*ml Primocin (Invitrogen # ant-pm-05). mESC maintenance media was supplemented with 1,000 units/mL of Recombinant Mouse Leukemia Inhibitory Factor (Millipore Sigma # ESG1107). Media was changed every day.

Vitronectin coating was incubated for 15 min at a concentration of 0.5 µg*/*mL dissolved in 1X Phosphate-buffered saline (PBS) pH 7.4 (Thermo Fisher Scientific # 70011044). Dissociation and cell passages were done using ReLeSR passaging reagent (Stem Cell Technologies # 05872) according to the manufacturer’s instructions. Cell freezing was done in mFreSR cryopreservation medium (Stem Cell Technologies # 05855) according to the manufacturer’s instructions.

### Mouse cortical organoid generation

Mouse cortical organoids were grown as previously described by our group [57, 62]; with some modifications. To generate cortical organoids we single cell dissociated ESCs using TrypLE Express Enzyme (ThermoFisher Scientific #12604021) for 5 minutes at 37°C and re-aggregated in lipidure-coated 96-well V-bottom plates at a density of 3,000 cells per aggregate, in 100 µL of mESC maintenance media supplemented with Rho Kinase Inhibitor (Y-27632, 10 µM, Tocris # 1254) and 1,000 units/mL of Recombinant Mouse Leukemia Inhibitory Factor (Millipore Sigma # ESG1107) (Day −1).

After one day (Day 0), we replaced the medium with cortical differentiation medium containing Glasgow Minimum Essential Medium (Thermo Fisher Scientific # 11710035), 10% Knockout Serum Replacement (Thermo Fisher Scientific # 10828028), 0.1 mM MEM Non-Essential Amino Acids (Thermo Fisher Scientific # 11140050), 1 mM Sodium Pyruvate (Millipore Sigma # S8636), 2 mM Glutamax supplement (Thermo Fisher Scientific # 35050061), 0.1 mM 2-Mercaptoethanol (Millipore Sigma # M3148), and 0.05 mg*/*ml Primocin (Invitrogen # ant-pm-05). Cortical differentiation medium was supplemented with Rho Kinase Inhibitor (Y-27632, 20 µM # 1254), WNT inhibitor (IWR1-*ϵ*, 3 µM, Cayman Chemical # 13659), and TGF-Beta inhibitor (SB431542, Tocris # 1614, 5 µM). Days 0-5 media were changed every day.

On day 5, organoids were transferred to ultra-low adhesion plates (Millipore Sigma # CLS3471) where media was aspirated and replaced with fresh neuronal differentiation media. The plate with organoids was put on an orbital shaker at 60 revolutions per minute. Neuronal differentiation medium contained Dulbecco’s Modified Eagle Medium: Nutrient Mixture F-12 with GlutaMAX supplement (Thermo Fisher Scientific # 10565018), 1X N-2 Supplement (Thermo Fisher Scientific # 17502048), 1X Chemically Defined Lipid Concentrate (Thermo Fisher Scientific # 11905031) and 0.05 mg*/*ml Primocin (Invitrogen # ant-pm-05). Organoids were grown under 5% CO2 conditions. The medium was changed every 2-3 days.

On day 14 and onward, we transferred the organoids to neuronal maturation media containing BrainPhys Neuronal Medium (Stem Cell Technologies # 05790), 1X N-2 Supplement, 1X Chemically Defined Lipid Concentrate (Thermo Fisher Scientific # 11905031), 1X B-27 Supplement (Thermo Fisher Scientific # 17504044), 0.05 mg*/*ml Primocin (Invitrogen # ant-pm-05) and 0.5% v/v Matrigel Growth Factor Reduced (GFR) Basement Membrane Matrix, LDEV-free.

### Immunohistochemistry and confocal imaging

Organoids were collected and fixed in room temperature 4% Paraformaldehyde (PFA) (Thermo Fisher Scientific # 28908) and cryopreserved in 30% Sucrose (Millipore Sigma # S8501). They were then embedded in a solution containing 50% of Tissue-Tek O.C.T. Compound (Sakura # 4583) and 50% of 30% sucrose dissolved in 1X Phosphate-buffered saline (PBS) pH 7.4 (Thermo Fisher Scientific # 70011044). Organoids were then sectioned to 20 µm using a cryostat (Leica Biosystems # CM3050) directly onto glass slides. After 2 washes of 5 minutes in 1X PBS and 1 wash in deionized water (Chem world #CW-DW-2G), the sections were incubated in blocking solution 5% v/v donkey serum (Millipore Sigma # D9663), and 0.1% Triton X-100 (Millipore Sigma # X100) for 1 hour. The sections were then incubated in primary antibodies overnight at 4°C. They were then washed 3 times for 10 minutes in PBS and incubated in secondary antibodies diluted in blocking solution for 90 minutes at room temperature. They were then washed 3 times for 10 minutes in PBS and coverslipped with Fluoromount-G Mounting Medium (Thermo Fisher Scientific # 00-4958-02).

Primary antibodies used were: mouse anti Gfap (Thermo Fisher Scientific #G6171, RRID: AB_1840893, 1:100); rabbit anti Map2 (Proteintech #17490-1-AP, RRID: AB_2137880, 1:2000); mouse anti Sst (Santa Cruz Biotechnology #sc-55565, RRID: AB_831726, 1:100); mouse anti Satb2 (Abcam #ab51502, RRID: AB_882455, 1:100); rabbit anti Tbr1 (Millipore Sigma #AB10554, RRID: AB_10806888, 1:500); mouse anti Pax6 (BD Biosciences #561462, RRID: AB_10715442, 1:100).

Secondary antibodies were of the Alexa series (Thermo Fisher Scientific), used at a concentration of 1:750. Nuclear counterstain was performed using 300 nM DAPI (4’,6-Diamidino-2-Phenylindole, Dihydrochloride) (Thermo Fisher # D1306).

Imaging was done using an inverted confocal microscope (Zeiss 880) and Zen Blue software (Zeiss). Images were processed using Zen Black (Zeiss) and ImageJ software (NIH).

## Experiment Phases

### Record Phase

In order to rapidly determine the location and footprint of the neurons to use for the experiment, we initially perform a spontaneous recording. This captures spontaneous neural activity, and does rapid, automatic analysis in order to locate neurons based on the quantity and magnitude of action potentials above a threshold. The spike detection threshold was chosen in the MaxLab Live software (MaxWell Biosystems) as 5 times the root-mean-square amplitude of the signal of the channel. (To ensure that the most relevant units are selected, this is more conservative than the threshold used for later phases.)

Channels with activity were then combined into putative neural unit footprints by taking a spike triggered average of electrical signal in all other channels. Channels whose spike-triggered average had peak-to-peak variation above an empirically chosen threshold were considered part of the same footprint; this forms an ad-hoc test for channels which are significantly likely to demonstrate activity at the same time. Redundant footprints were eliminated by finding pairs which shared at least 50% of their spike times, and retaining the one with larger peak-to-peak signal amplitude in its own spike-triggered average.

From these footprints, up to 32 putative units were selected for stimulation based on a custom metric *η*, designed to identify the electrodes where spikes could be detected most consistently by combining the firing rate and mean spike amplitude *µ*_amp_ as follows:

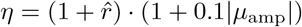

The normalized log spike rate 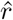 is defined with respect to the minimum and maximum spike rates across all electrodes as follows:

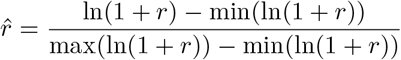

### Stimulation Phase

After identifying putative neural units, we characterize their stimulus-response properties by delivering biphasic pulses to each unit and recording the network’s response. Previous studies have identified multiple different response timescales [63], so we consider two separate connectivity metrics: the first-order connectivity *C*_1_, and the multi-order connectivity *C*_*m*_.

For a window of stimulus-relative time, we define a response tensor indexed by the stimulus repetition *k*, the stimulus electrode *i*, the recording channel *j*.

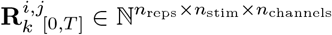

where *n*_reps_ is the number of stimulus repetitions (50), *n*_stim_ is the number of stimulation electrodes corresponding to putative neurons, and *n*_channels_ is the number of recording channels specifically under putative neurons. The value of the tensor is the number of spikes in an interval of time relative to each stimulus, with spikes identified by thresholding at 3 standard deviations.

First-order connectivity *C*_1_ is the observed probability of evoking at least one spike within 10ms:

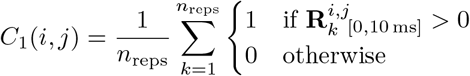

Multi-order connectivity *C*_*m*_ is the average number of spikes observed in a longer window:

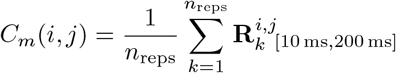

To identify network-wide bursts of the units, we compute the total spike count across all channels for each stimulation. A burst is detected when this count exceeds the median count plus three median absolute deviations (MAD). Bursts are excluded from the multi-order connectivity calculation to focus on specific neural pathways rather than larger network activations.

These metrics enable identification of both direct and network-mediated connections between neural units.

### Cartpole Environment

Cartpole (also known as inverted pendulum) is a traditional control problem which tests an agent’s ability to balance an unstable dynamical system. The task is typically restricted to 2D, involving a cart that can move horizontally balancing a pole vertically that is free to rotate around the attachment joint. The goal is to keep the pole upright by applying horizontal forces to the cart. This benchmark is valuable for its simple formulation yet inherent instability, real-world parallels, and widespread use in both control theory and reinforcement learning [46, 47].

The system state is fully described by four variables: cart position *x*, cart velocity 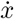, pole angle *θ*, and angular velocity 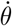. The agent can apply a force *F* ∈ {−10N, 10N}to the cart at each timestep. The full equations of motion describing the nonlinear dynamics can be found in [45]. Episodes begin with small random perturbations to *θ* and 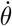 and proceed in discrete timesteps until |*θ*| *>* 16^*°*^, at which point the pole is considered to be in an unrecoverable state. We chose to remove traditional horizontal position bounds to focus solely on the pole balancing aspect of the task.

Each timestep involves decoding spike rates from the culture to determine the applied force, updating the virtual environment state with said force, and adjusting the encoding stimulation rates corresponding to the current virtual state.

### Encoding and Decoding Information

Throughout this study, we use rate coding for both input/encoding and output/decoding signals, which provides sufficient information transmission, especially when focusing on small numbers of neurons [10, 64, 65]. The virtual state is encoded to the culture through two input neurons receiving stimulation frequencies determined by the pole angle *θ*:

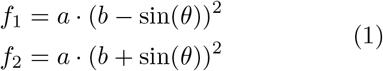

where *a* = 7 and *b* = 0.15 are scaling factors that maintain stimulation frequencies in a biologically relevant range (both neurons receive ≈ 1.1 Hz when the pole is vertical, with frequencies diverging to ≈0.8 Hz and ≈8.9 Hz at terminal angles) while still being able to represent the sign of the angle. This functional form was chosen because the sinusoidal encoding ensures that the neurons receive opposing signals based on the pole’s deviation from vertical, while the exponent enables the frequency to change faster as it nears the terminal angle.

For decoding signal from the culture, spikes are detected by a threshold of 3 standard deviations. Each output neuron’s activity is then converted to a smoothed firing rate by an exponential moving average:

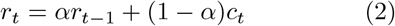

where *r*_*t*_ is the smoothed firing rate at time *t, c*_*t*_ is the raw spike count in the current window, and *α* = 0.2 balances immediate responses with firing rate stability.

The force *F* applied to the cart is determined by the difference between the two output units’ smoothed firing rates, which are clipped to the range from −1 to 1 and then multiplied by 10 N.

### Real-time Implementation

The closed-loop system operates through discrete timesteps involving three key components: the organoid’s neural activity, the simulated environment, and the training signals. Each timestep consists of:

1. Read Phase (200 ms):
  - Process raw signals using SALPA (Subtraction of Artifacts by Local Polynomial Approximation) [66] for artifact removal
  - Detect spikes via thresholding at 3 standard deviations, or set threshold above noise
  - Monitor motor neuron activity via smoothed firing rates *r*_*t*_ (Eq. 2)
2. Environment Update:
  - Decode motor signals *r*_1,*t*_ and *r*_2,*t*_ to output force *F*
  - Update cartpole state according to physics
  - Encode new state to stimulation frequencies via Eq. 1
3. Training Phase (200 ms, conditional):
  - Skip unless episode completed
  - Deliver training patterns if 5-episode mean performance below 20-episode mean
  - For adaptive protocol (see below), update value estimates based on performance

Each phase requires precise timing to maintain real-time control. A longer read phase yields reliable spike counts and smoother actions, whereas a shorter read phase leads to more responsive, yet noisy control. Experiments used either 200 ms or 300 ms read phases. For direct comparison, performance metrics from 200 ms experiments were scaled by a factor of 1.5 to normalize to a 300 ms time base.

### Training Signal Implementation

Three experimental conditions were tested: a null case without stimulation, and two stimulation paradigms, random and adaptive, with signals administered conditionally based on performance metrics, ether to randomly chosen training neurons or adaptively selected training neurons. Specifically, stimulation occurred when the 5-episode mean performance fell below the 20-episode moving average, allowing the system to focus on larger-scale adaptations while filtering out the impact of individual poor episodes. The pool from which individual training neurons and corresponding electrodes (“training electrodes”) were selected to receive feedback stimulation (*n* = 8–15) was created in the Record phase based on the activity metric described in Methods: Record Phase.

The random paradigm delivered sequential biphasic pulses (5 ms inter-pulse interval) through 5 training electrodes selected randomly without replacement. Each stimulation epoch consisted of one complete pulse sequence delivered at 10 Hz for 300 ms.

In contrast to the random paradigm, the adaptive paradigm utilized a weighted sampling approach for paired-pulse patterns (5 ms inter-pulse interval). Each electrode *i* had a value estimate *V*_*i*_ updated at the end of each episode via:

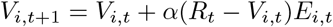

where *R*_*t*_ denotes time balanced (our performance metric, episode duration), *α* (0.3) is the learning rate, and *E*_*i*,*t*_ represents the eligibility trace:

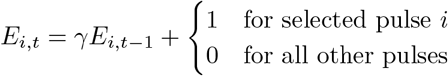

where *γ* (0.3) is the decay factor that diminishes the influence of past selections. The eligibility trace enables temporal credit assignment by storing a decaying record of recent stimulation patterns. weighted by their previous success in improving performance.

Value estimates were not allowed to decrease between the minimum possible episode reward of 10. Selection probabilities were computed as:

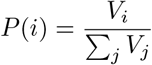

Training patterns were sampled according to these probabilities and delivered as paired pulses at 10 Hz for 300 ms.

### Statistical Analysis

Prior to statistical testing, we assessed normality of performance distributions using Shapiro-Wilk tests, which revealed significant deviations from normality across all conditions (*p <* 0.001). Given this non-normal distribution, we employed the non-parametric Mann-Whitney U test for pairwise comparisons between conditions (null, random, adaptive, and continuous adaptive), with Holm-Bonferroni correction for multiple comparisons.

Performance thresholds were determined based on the 99th percentile of simulated random controllers (see Supplementary Methods). A cycle was considered “proficient” if its 90th percentile time balanced exceeded this threshold. All statistical tests used an alpha level of 0.05, with *p*-values reported where appropriate.

## Acknowledgments

This work was supported by the Schmidt Futures Foundation SF 857 and the National Human Genome Research Institute under Award number 1RM1HG011543 (D.H. and M.T.), the National Science Foundation under award number NSF 2034037 (M.T), and NSF 2134955 (M.T. and D.H), and the National Institute of Mental Health under award number 1U24MH132628 (M.A.M.-R. and D.H.). We are thankful to the Pacific Research Platform, supported by the National Science Foundation (NRP) under award numbers CNS-1730158, ACI-1540112, ACI-1541349, and OAC-1826967, the University of California Office of the President, and the University of California San Diego’s California Institute for Telecommunications and Information Technology/QualcommInstitute.

## Funding

- Schmidt Futures Foundation SF 857
- National Human Genome Research Institute grant 1RM1HG011543 (DH, MT)
- National Science Foundation grant 2034037 (MT)
- National Science Foundation grant 2134955 (MT, DH)
- National Institute of Mental Health grant 1U24MH132628 (MAM-R, DH)

## Competing interests

K.V. is a co-founder and D.H., M.T. are advisory board members of Open Culture Science, Inc., a company that may be affected by the research reported in the enclosed paper. All other authors declare no competing interests.

## Notes

https://braingeneers.github.io/braindance/

https://github.com/braingeneers/braindance

